# Amygdala stimulation transforms short-term memory into remote memory by persistent activation of atypical PKC in the anterior cingulate cortex

**DOI:** 10.1101/2024.09.17.613040

**Authors:** William Almaguer-Melian, Daymara Mercerón-Martínez, Laura Alacán-Ricardo, Arturo Bejerano Pina, Changchi Hsieh, Jorge A. Bergado-Rosado, Todd Charlton Sacktor

## Abstract

Although many studies have addressed the role of the amygdala in modulating long-term memory, it is not known whether weak training plus amygdala stimulation can transform a short-term memory into a remote memory. Object place recognition (OPR) memory after strong training remains hippocampus-dependent through the persistent action of PKMζ for at least 6 days, but it is unknown whether weak training plus amygdala stimulation can transform short-term memory into an even longer memory, and whether such memory is stored through more persistent action of PKMζ in hippocampus. We trained rats to acquire OPR and 15 min or 5 h later induced a brief pattern of electrical stimulation in basolateral amygdala (BLA). Our results reveal that a short-term memory lasting < 4 h can be converted into remote memory lasting at least 3 weeks if the BLA is activated 15 min, but not 5 h after learning. To examine how this remote memory is maintained, we injected ZIP, an inhibitor of atypical PKCs (aPKCs), PKMζ and PKCι/λ, into either hippocampal CA1, dentate gyrus (DG), or anterior cingulate cortex (ACC). Our data reveal amygdala stimulation produces consolidation into remote memory, not by persistent aPKC activation and capture by synaptic tagging processes in the hippocampal formation, but in ACC. Our data establish a powerful modulating role of the BLA in forming remote memory and open a path in the search for neurological restoration of memory, based on enhancing synaptic plasticity in aging or neurodegenerative disorders such as Alzheimer’s disease.

## Introduction

Learning and memory constitutes the basis by which animals build internal models of the world in the domains of space and time, allowing useful predictions founded upon stored experience.

Understanding the mechanisms of memory is one of the fundamental challenges of neuroscience; in particular, how, of all the information that our nervous system continuously processes, only some is selected for permanent storage. There is extensive literature showing that the basolateral amygdala (BLA) is a critical region for memory consolidation (McGaugh JL, 2002; McGaugh JL, 2015; Parent MB *et al*., 1995; Roozendaal B and McGaugh JL, 1997). Pharmacological studies have shown that post-training BLA interference disrupts the consolidation of memories (Hatfield T and McGaugh JL, 1999; Nedaei SE *et al*., 2016; Packard MG *et al*., 1994; Sardari M *et al*., 2015), whereas post-training intra-BLA administration of norepinephrine promotes memory consolidation (LaLumiere RT *et al*., 2003; Roesler R *et al*., 2021). Moreover, β-adrenergic antagonists administered in BLA prevents the memory reinforcement produced by systemic administration of epinephrine, glucocorticoid agonists, the opioid antagonist naltrexone, or ketamine (Campolongo P *et al*., 2009; Liang KC *et al*., 1986; Morena M *et al*., 2021; Quirarte GL *et al*., 1997). In addition, noradrenergic activation of the BLA enhances the consolidation of object recognition memory, suppressing anterior insular cortex activity (Chen Y *et al*., 2018; Chen YF *et al*., 2022). More recently, optogenetic stimulation of BLA projections to the medial entorhinal cortex immediately after memory acquisition was found to enhance spatial and contextual memory retention (Wahlstrom KL *et al*., 2018). Studies in humans furthermore showed that electrical stimulation of the amygdala leads to improved memory (Inman CS *et al*., 2018). Interestingly, we have repeatedly observed that basolateral amygdala stimulation can improve spatial learning skills in rats with a profound cognitive deficits produced by lesions of the fimbria fornix, suggesting that similar procedures could contribute to the recovery of memory capacities in humans (Almaguer-Melian W *et al*., 2005; Mercerón-Martínez D *et al*., 2018; Mercerón-Martínez D *et al*., 2020; Merceron-Martinez D *et al*., 2016; Mercerón-Martínez D *et al*., 2016; Mercerón-Martínez D *et al*., 2022; Mercerón-Martínez D *et al*., 2013).

In addition to BLA regulating memory consolidation, studies of synaptic plasticity such as long-term potentiation (LTP), a cellular model of memory (Bailey CH *et al*., 2015; Bliss TV and Lomo T, 1973; Matthies H, 1989), suggest underlying mechanisms for how amygdala activity can modulate memory. These studies have shown that amygdala stimulation can modulate LTP in the dentate gyrus (Akirav I and Richter-Levin G, 1999; Akirav I and Richter-Levin G, 1999; Ikegaya Y *et al*., 1994; Ikegaya Y *et al*., 1995; Ikegaya Y *et al*., 1995). Likewise, emotional-motivational behavioral stimulation can prolong a short-lasting early-LTP (E-LTP) into a long-lasting late LTP (L-LTP) (Almaguer-Melian W *et al*., 2010; Seidenbecher T *et al*., 1997). Similarly, direct stimulation of the basolateral amygdala within a time window close to that of LTP induction, prolongs early-LTP into late-LTP (Frey S *et al*., 2001). Conversely, transient or permanent inactivation of the BLA abolishes the behavioral reinforcing effects on LTP (Almaguer-Melian W *et al*., 2003). The neural pathways involved in the BLA reinforcement of LTP include adrenergic terminals from the locus coeruleus and cholinergic afferents from the medial septum (Bergado JA *et al*., 2007). More recently, we have provided data showing that BLA stimulation for 15 min daily for 4 days after water maze training promotes recovery of spatial memory and LTP that was impaired by fimbria-fornix injury, suggesting a possible functional relationship between the recovery of synaptic plasticity and the recovery of memory (Mercerón-Martínez D, *et al*., 2022).

Place recognition memory after strong training is hippocampus-dependent through the persistent action of PKMζ for at least 6 days (Hardt O *et al*., 2010), but it is not known whether weak training plus amygdala stimulation can transform a short-term memory (□4 h) into an even longer remote memory lasting weeks, and whether such a memory might be stored through even more persistent activation of PKMζ in hippocampus. Memory and LTP share molecular mechanisms that, in a temporally sequenced manner, sustain their expression through different phases (Reymann K *et al*., 1988). Intermediate and long-term phases depend on multiple protein kinases and protein synthesis during its initial consolidation period (Krug M *et al*., 1984). PKMζ, an unusual, persistently active atypical PKC (aPKC) isoform (Sacktor TC and Hell JW, 2017), together with the redundant, closely related aPKC, PKC□/ □, have been proposed as fundamental players in long-term memory storage beyond this initial consolidation (Patel H and Zamani R, 2021; Tsokas P *et al*., 2016). Therefore, we were interested in exploring the role of persistently active aPKC in long-term remote memory. To this purpose, we applied the aPKC inhibitor ZIP (Sacktor TC and Fenton AA, 2012) to trained animals to ascertain whether BLA stimulation can maintain remote memory by persistently activating PKMζ or PKC□/ □, either in the hippocampus or in cortical regions such as the anterior insular cortex (ACC). Our data suggest that BLA stimulation within the initial modulation window of neural plasticity (Frey U and Morris RG, 1997; Frey U and Morris RG, 1998) induces remote memory formation by the activation and capture of aPKC in synapses tagged by the experience-related activity in ACC. These results show that concurrent amygdala activation can convert an incidental short-term memory of daily experience (□4h) into a remote memory that could potentially last a lifetime in neocortex, opening new possibilities for its use to improve memory capacity in memory disorders such as Alzheimer’s disease.

## Materials and Methods

### Animals

We used 147 eight-week-old male Wistar rats weighing 270 to 300 g at the beginning of the experiment. Animals were provided by the Cuban National Center for Laboratory Animals (CENPALAB) and maintained in translucent plastic cages (5 animals per cage) under controlled environmental conditions (23 °C, constant humidity, 12-h light-dark cycles) with free access to food and water throughout the experiment.

### Ethics

All rats were handled and maintained according to the international ethic norms for the use of laboratory animals, and abide by the National Institute of Health Guide for the Care and Use of Laboratory Animals (NIH Publications No. 80-23) revised 1996, the UK Animals (Scientific Procedures) Act 1986 and associated guidelines, and the European Communities Council Directive of 24 November 1986 (86/609/EEC). We also comply with the Cuban regulations published by CENPALAB and the internal regulations of the International Center for Neurological Restoration (CIREN). All efforts were made to reduce pain and discomfort. We have formed experimental groups with the smallest number of animals without affecting the methodological robustness of the experiments. Three days before beginning the behavioral studies, the animals were gently handled by the experimenter for about 2 h to habituate them and reduce handling stress.

### Chronic implantation of electrodes and intra-cerebral injection cannula

Electrodes and injection cannula were stereotactically implanted under anesthesia by intraperitoneal injection of ketamine (50 mg/10 ml), diazepam (10 mg/1 ml), and atropine (0.5 mg/1 ml) in a volume of 1 mL/100 g body weight. We bilaterally implanted the cannula in dentate gyrus (DG; anteroposterior [AP] = -3.8 mm, mediolateral [ML] = ±2.0 mm, and dorsoventral [DV] = -3.5 mm from bregma), cornu ammonis 1 (CA1; AP = −4.0; ML = ±4.0, and DV = −3.0 mm from bregma) or anterior cingulate cortex (ACC; AP = +2.7; ML = ±0.6, and DV = -2.7), according to experimental group. After the cannula were fixed with dental acrylic cement, we proceeded to bilaterally implant BLA electrodes (AP = –3.1 mm, ML = ±4.7 mm, and DV = -8.5 mm) (Paxinos G and Watson C, 1998). We attached 3 miniscrews to strengthen the preparation, and the electrodes were connected to a socket and fixed with dental acrylic cement.

### Object Placement Recognition memory test

To test our working hypothesis we used Object Placement Recognition (OPR), a one-trial spatial memory task, based on the spontaneous activity of the animals to preferentially explore novelty within a familiar environment. The behavioral task was completed in three steps. On the first day, habituation was performed in an open field (50 x 50 cm) illuminated by a 40 W light bulb placed 1 m above the floor, and on one of its walls a spatial cue, consisting of a letter-type sheet with a bold black capital A, 650 dpi, was fixed. Animals were placed in the center of the open field, facing the spatial reference and allowed to freely explore the arena for 5 min. Twenty-four hours later memory was acquired by placing two similar objects in the open field. The animals were free to explore both objects for 3 min. The third step was the memory retention test, which was performed 2 h, 4 h, 24 h, or 21 days after acquisition. For the retention test, one of the objects (left block) was moved to a new location, and the animals were allowed to explore freely for 3 min. We measured exploration time for an object as the time in which the rat stayed with its head close to the front of the object or with its forelimbs on the object. In both memory acquisition and the memory retention test, the exploration time of each individual object was measured and expressed as the percentage of the total exploration time of the two objects. All behavioral studies were conducted between 09:00-12:30 h to reduce circadian influences.

### BLA stimulation

The stimulation was applied simultaneously to both BLA regions and consisted of 3 trains of 15 impulses at 200 Hz, as previously described (Frey S, *et al*., 2001). Each stimulus was 0.2 ms in duration, and trains were delivered with 10 s intertrain intervals. We applied this tetanus 15 min or 5 h after memory acquisition. The stimulation intensity was 400 μA to each side.

### Experimental groups

The study design consisted of three experiments. Experiment I was to establish baseline values of retention without BLA stimulation. Experiment II was to study the effect of BLA stimulation on the formation of long-term and remote memory. Experiment III was to examine whether the maintenance of remote memory (21 days) facilitated by amygdala stimulation depends on persistent aPKC action. To test this we injected ZIP (10 nmol in 1 □L in each side; Tocris Bioscience, UK) or Vehicle (NaCl, 0.9%, pH 7.0) on day 19 in DG, CA1, or ACC.

#### Short- and long-term memory

1. Control 2 h: rats performed OPR memory test 2 h post-acquisition trial, n = 10
2. Control 4 h: rats performed OPR memory test 4 h post-acquisition trial, n = 13
3. Control 24 h: rats performed OPR memory test 24 h post-acquisition trial, n = 24

#### Amygdala stimulation facilitating long-term memory to remote memory

4. BLA-15 min + 24 h: rats received amygdala stimulation 15 min post-acquisition and performed OPR memory test 24 h post-acquisition trial, n = 7
5. BLA-5 h + 24 h: rats received amygdala stimulation 5 h post-acquisition and performed OPR test 24 h post-acquisition trial, n = 11
6. Control+21days: rats performed OPR memory test 21 days post-acquisition trial, n = 10
7. BLA-15 min + 21 days: rats received amygdala stimulation 15 min post-acquisition and performed OPR memory test 21 days post-acquisition trial, n= 10

#### Mapping aPKC-dependent remote memory storage by amygdala stimulation

8. Control DG + 21 days: rats received OPR memory testing 21 days post-acquisition, n = 14
9. BLA DG-NaCl: rats received amygdala stimulation 15 min post-acquisition, NaCl injections into DG on day 19 post-acquisition, and OPR memory testing 21 days post-acquisition, n = 9
10. BLA DG-ZIP: rats received amygdala stimulation 15 min post-acquisition, ZIP injections into DG on day 19 post-acquisition, and OPR memory testing 21 days post-acquisition, n = 8
11. Control CA1+ 21 days: rats received OPR memory testing 21 days post-acquisition, n= 10
12. BLA CA1-NaCl: rats received amygdala stimulation 15 min post-acquisition, NaCl injections into CA1 on day 19 post-acquisition, and OPR memory testing 21 days post-acquisition, n = 11
13. BLA CA1-ZIP: rats received amygdala stimulation 15 min post-acquisition, injections into CA1 on day 19 post-acquisition, and OPR memory testing 21 days post-acquisition, n = 9
14. BLA+ACC-NaCl: rats received amygdala stimulation 15 min post-acquisition, NaCl injections into AAC on day 19 post-acquisition, and OPR memory testing 21 days post-acquisition, n = 7
15. BLA+ACC-ZIP: rats received amygdala stimulation 15 min post-acquisition, injections into AAC on day 19 post-acquisition, and OPR memory testing 21 days post-acquisition, n = 6

### Statistical analysis

We first confirmed the normal distribution of the data using the Kolmogorov Smirnov test, and variance homogeneity by the Barlett’s test. For statistical analysis of data during habituation we used ANOVA for repeated measures. To compare results among more than two groups, a one-way ANOVA was performed, using the Tukey’s test for the *post hoc* analysis. Statistical significance was defined as p < 0.05. In the graphs that appear in the figures, letters were placed to specify the statistically significant differences between the exploration time during the acquisition and the memory test, and between the experimental groups. Equal letters denote that there are no statistically significant differences, whereas different letters denote statistically significant differences.

## Results

### Short-term memory for OPR

Animals were first habituated to an empty arena (Fig. 1A), in which the number of visits to the borders of the box rapidly decreases (repeated measures ANOVA, main effect of time is significant, *F*_4, 84_ = 243.22, *p* < 0.000001, followed by the *post hoc* test of Tukey HSD; different letters denote significant differences; same letters denote no significant differences). As in subsequent experiments, the number of visits is indistinguishable between the groups that were randomly selected to be trained the next day (repeated measures ANOVA, main effect of group, *F*_1, 21_ = 2.14, *p* = 0.28).

**Fig. 1.**
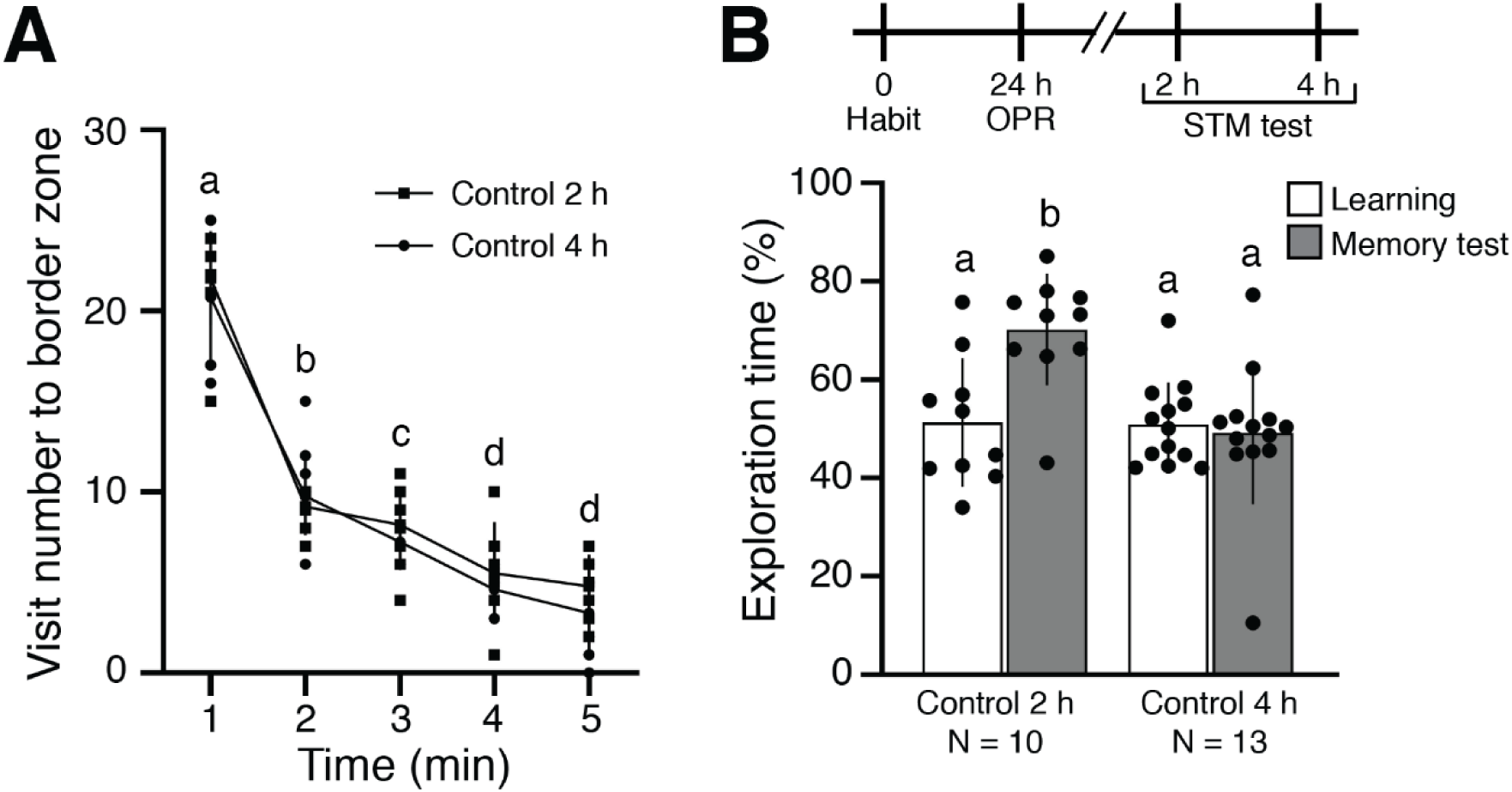
Behavioral performance in OPR during short-term memory retrieval. (A) Habituation of 5-min exposures to open field show no difference between 2 h and 4 h groups. (B) OPR retrieval test during short-term memory shows retention at 2 h but not 4 h after learning. Above, schematic of experimental procedures. Below, means and standard error of the mean. Different letters denote statistical differences, and same letters denote no statistical differences.

Twenty-four hours after habituation, animals were returned to the arena, which now contained two identical objects. Fig. 1B shows that whereas animals have no initial preference for one object or the other, they explore the object that had been moved to a different location when the retention test is carried out 2 h after training, but not when testing is delayed to 4 h (Fig. 1B, repeated measures ANOVA, main effect of group, *F*_1, 42_ = 7.11, *p* = 0.00057, interaction of group and learning-memory test, *F*_1, 42_ = 8.23, *p* = 0.00064, followed by *post hoc* Tukey HSD test; different letters mean statistical differences; same letters denote no significant differences).

### Amygdala stimulation facilitates long-term and remote OPR memory formation

We next examined whether amygdala stimulation after weak training facilitates the formation of long-term memory. Prior to training, habituation of animals is similar without any group difference (Fig. 2A: repeated measures ANOVA, main effect of time, *F*_4, 108_ = 127.32, *p* < 0.000001; group, *F*_2, 27_ = 0.18, *p* = 0.84, followed by *post hoc* test of Tukey HSD; different letters denote significant differences; same letters denote no significant differences). After short-term memory training, we stimulated the BLA and tested 24 h later. When the BLA was stimulated 15 minutes after training, retention of object placement memory was significantly longer than that of control, non-stimulated animals, or animals that receive BLA stimulation 5 hours after training (Fig. 2B, repeated measures ANOVA, main effect of group, *F*_2, 54_ = 8.99, *p* = 0.000003, interaction of group and learning-memory test, *F*_2, 54_ = 8.42, *p* = 0.00066, followed by *post hoc* test of Tukey HSD; different letters denote significant differences; same letters denote no significant differences). Control non-stimulated animals show no retention. Thus, BLA activation prolongs a short-term memory trace, but only when it occurs within a time window close to the training.

**Fig. 2.**
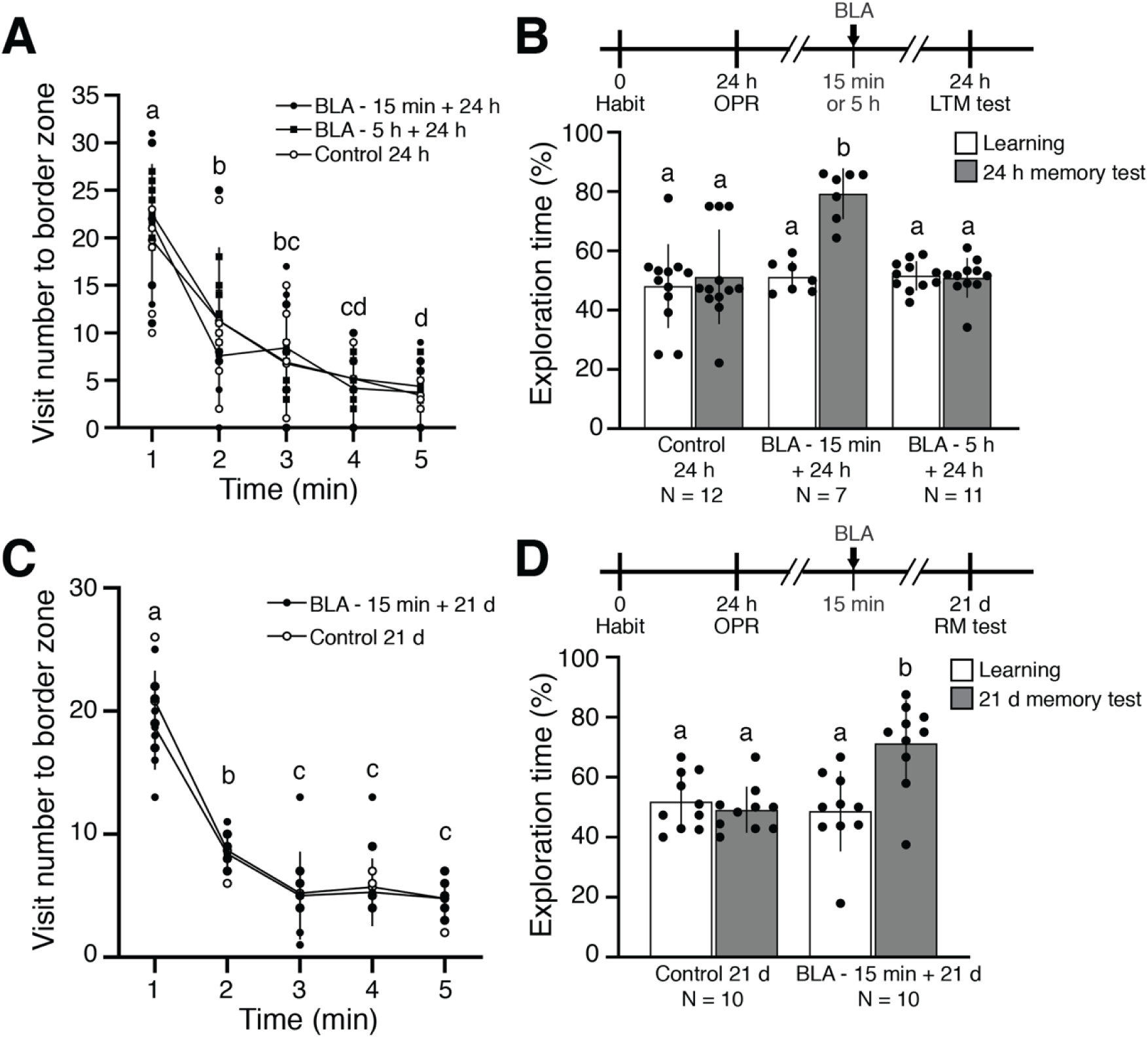
Behavioral performance in the OPR task during long-term and remote memory retrieval facilitated by amygdala stimulation. (A) Habituation of 5-min exposures to open field show no difference between BLA stimulated at 15 min, BLA stimulated at 5 h, and control groups. (B) Effect of time-dependent BLA stimulation on prolonging memory duration. Above, schematic of the experiment procedures. Below, means and standard error of the mean. (C) Habituation of 5 min exposures to open field. (D) OPR memory retention test shows BLA stimulation facilitates remote memory formation. Above, schematic of experimental procedures. Below, means and standard error of the mean. Different letters denote statistical differences, and same letters denote no statistical differences.

We then assessed whether amygdala stimulation also promotes remote memory formation lasting weeks. Group behaviors during the habituation task were not different (Fig. 2C: repeated measures ANOVA, main effect of time, *F*_4, 72_ = 142.73, *p* < 0.000001; group, *F*_1, 18_ = 1.78, *p* = 0.29, followed by the *post hoc* test of Tukey HSD; different letters denote significant differences; same letters denote no significant differences). Stimulation of the amygdala 15 min after training causes the exploration time of the object in the new position to be greater than that of the object placed in the familiar position 21 days after training (Fig. 2D: repeated measures ANOVA, main effect of group, *F*_1, 36_ = 8.60, *p* = 0.00019, interaction of group and learning-memory test, *F*_1, 36_ = 11.87, *p* = 0.0015, followed by *post hoc* test of Tukey HSD; different letters denote significant differences; same letters denote no significant differences).

### Remote OPR memory storage facilitated by amygdala stimulation depends on persistent aPKC action in ACC

We then examined the location of long-term memory storage for BLA stimulation-facilitated OPR. Strong OPR training produces a memory that remains hippocampus-dependent and sensitive to ZIP for at least 6 days, and thereafter fades (Hardt O, Migues PV, Hastings M, Wong J and Nader K, 2010). BLA stimulation might make this hippocampus-dependent memory more persistent.

Alternatively, memories can be initially stored in hippocampus and then become dependent on extrahippocampal regions during remote memory storage, and BLA stimulation might induce this consolidation at the systems level. To distinguish between these possibilities, we inhibited PKMζ in three memory-relevant regions: DG, CA1, and ACC. Figs. 3A, C, and E show habituation without group difference (Fig. 3A: repeated measures ANOVA, main effect of time, *F*_4, 124_ = 368.58, *p* < 0.000001, group, *F*_2, 31_ = 1.57, *p* = 0.22; Fig. 3C, time, *F*_4, 108_ = 70.03, *p* < 0.000001, group, *F*_2, 27_ = 0.27, *p* = 0.77; Fig. 3E: time, *F*_4, 44_ = 8, 46, *p* < 0.000001, group, *F*_1, 11_ = 1.02, *p* = 0.33, followed by the *post hoc* test of Tukey HSD; different letters denote statistical differences; same letters denote no significant differences). We injected ZIP 19 days after training, and then tested retention 2 days later, a duration that allows for elimination of ZIP in order to test the inhibitor’s effect on memory maintenance (Pastalkova E *et al*., 2006). The results reveal that blocking aPKC in the DG or CA1 has no disruptive effect on remote memory retention after training facilitated by BLA stimulation (Figs. 3B, D; Fig 3B repeated measures ANOVA, main effect of group, *F*_2, 62_ = 6.66, *p* = 0.000056, interaction of group and learning-memory test, *F*_2, 52_ = 4.59, *p* = 0.014; Fig. 3D: repeated measures ANOVA, main effect of group, *F*_2, 54_ = 9.64, *p* = 0.000001, interaction of group and learning-memory test, *F*_2, 54_ = 6.68, *p* = 0.0026, followed by the *post hoc* test of Tukey HSD; different letters denote statistical differences; same letters denote no significant differences). In striking contrast, ZIP completely blocks memory when applied in the ACC (Fig. 3F, repeated measures ANOVA, main effect of group, *F*_1, 22_ = 38.97, *p* < 0.000001, interaction of group and learning-memory test, *F*_1, 22_ = 26.98, *p* = 0.000033, followed by the *post hoc* test of Tukey HSD; different letters denote statistical differences; same letters denote no significant differences), indicating persistent aPKC activity in this brain region maintains remote OPR memory.

**Fig. 3.**
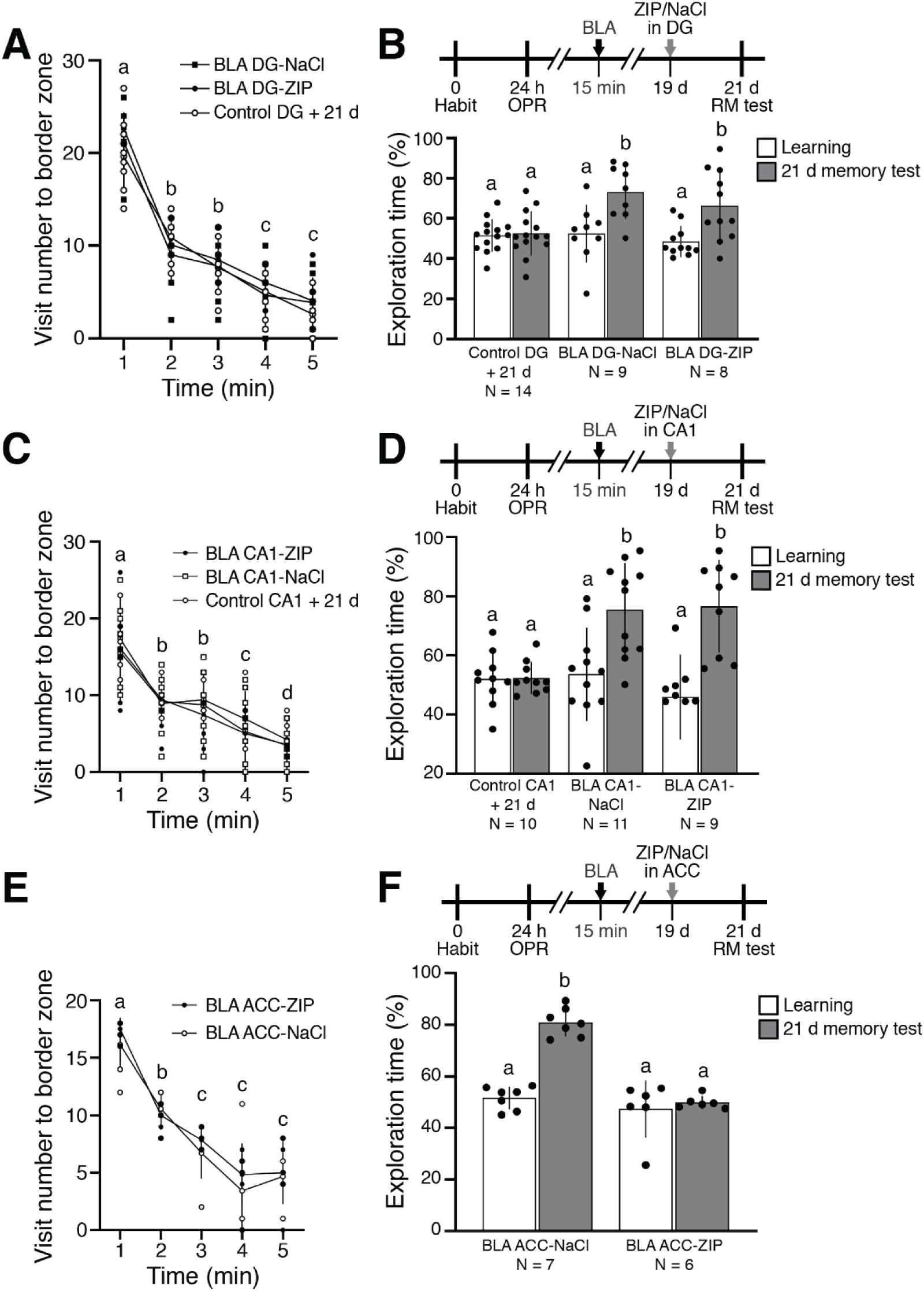
ZIP administration in DG, CA1, or ACC on behavioral performance in the OPR task during remote memory facilitated by amygdala stimulation reveals only ACC administration disrupts remote memory maintenance. (A, C, E) Habituation for 5-min exposures to open field. (B, D, F) Effects of intracranial DG, CA1, or ACC ZIP administration on BLA-stimulated remote memory. Above, schematic of experimental procedures. Below, means and standard error mean. (B) DG shows no effect. (D) CA1 shows no effect (F) ACC shows disruption of remote memory. Different letters denote statistical differences, and same letters denote no statistical differences.

## Discussion

In this work we obtained evidence that activation of the amygdala close to the time of memory acquisition produces a consolidation at the systems level through the processes of synaptic tagging and capture of aPKC in the anterior cingulate cortex, all of which contributes to establishing and maintaining a memory remote from experience. We found that BLA activation can facilitate not only the consolidation of short-term memory lasting less than 4 hours (Fig. 1B) into long-term memory lasting days (Fig. 2B), but also into remote memory lasting weeks (Fig. 2D, Fig. 3B, D, F). This consolidation occurs if the BLA is activated 15 min, but not 5 h, after acquisition (Fig. 1B).

Our data are in line with previous reports showing that the BLA is a critical region involved in memory consolidation (Campolongo P, *et al*., 2009; Chen Y, *et al*., 2018; Chen YF, *et al*., 2022; Hatfield T and McGaugh JL, 1999; LaLumiere RT, *et al*., 2003; Liang KC, *et al*., 1986; McGaugh JL, 2002; McGaugh JL, 2015; Morena M, *et al*., 2021; Nedaei SE, *et al*., 2016; Packard MG, *et al*., 1994; Parent MB, *et al*., 1995; Quirarte GL, *et al*., 1997; Roozendaal B and McGaugh JL, 1997; Sardari M, *et al*., 2015; Wahlstrom KL, *et al*., 2018), modulating synaptic plasticity processes such as LTP (Akirav I and Richter-Levin G, 1999; Akirav I and Richter-Levin G, 1999; Almaguer-Melian W, *et al*., 2010; Almaguer-Melian W, *et al*., 2003; Bergado JA, *et al*., 2007; Frey S, *et al*., 2001; Ikegaya Y, *et al*., 1994; Ikegaya Y, *et al*., 1995; Ikegaya Y, *et al*., 1995; Seidenbecher T, *et al*., 1997) that are maintained by the aPKC isoform, PKMζ (Ling DS *et al*., 2002; Pastalkova E *et al*., 2006; Tsokas P *et al*., 2024). To these findings, our results add a new complexity at the systems level to the amygdala’s modulatory role in encoding memory duration. Our evidence also indicates that remote memory facilitated by BLA activation is mediated by persistent aPKC activity in the ACC, because the aPKC-inhibitor ZIP abolishes memory when administered on day 19 post-acquisition in ACC, but not in CA1 or DG of the hippocampal formation. In addition, our results open a path to the search for neurological restoration based on synaptic plasticity of memory loss in aging and neurodegenerative diseases like Alzheimer’s disease.

There is general consensus that episodic memory initially depends on the hippocampus (recent memory) as it contributes to coordinating the activation of synapses widely distributed throughout the cortex that participate in memory encoding. Furthermore, successive reactivations of these memory traces coordinated by the hippocampus, contribute to increasing the efficiency of transmission between its cortical elements, thus creating a cortical representation of the experience that is less dependent on the hippocampus, known as remote memory (Dudai Y, 2004; Dudai Y *et al*., 2015; Hardt O and Nadel L, 2018; Kandel E *et al*., 2014; Squire LR *et al*., 2015). This transformation or stabilization of memory seems to imply that memory becomes more generalized, less precise in details (Hardt O and Nadel L, 2018). However, our data suggest that the spatial memory trace facilitated by amygdala activation retains a certain precision even at a remote moment in time, as the animals detect the spatial rearrangement produced by the change in position of one of the objects. In line with this result, previous findings from our laboratory show that animals in which the formation of remote place recognition memory was facilitated by administration of erythropoietin fail to detect the new spatial change if the spatial cue is removed, suggesting that even at this remote time period memory is spatial (Almaguer-Melian W *et al*., 2024). It has been suggested that latent engrams formed in contextual fear conditioning in the DG after systemic consolidation may again contribute to retrieval by providing rich contextual information. OPR, a non-reenforced learning task, is different than fear conditioning, but two considerations need to be taken into account. First, we stimulate the BLA, which in previous studies we have shown produces high c-Fos expression in both prefrontal cortex and DG cells (Mercerón-Martínez D, *et al*., 2020). These amygdala-activated DG cells could contribute to building a latent trace in the DG that can be reactivated at a remote time during memory evocation, providing vividness to the generalized cortical trace (Alam MJ *et al*., 2018; Tonegawa S *et al*., 2018). Second, in the place recognition memory task, unlike other tests such as contextual fear conditioning, a change or novelty is introduced in the spatial structure by changing one of the objects to a new position. This novelty can contribute to activating, among many other structures, the hippocampus, particularly the DG (Terranova JI *et al*., 2019; Tonegawa S, *et al*., 2018), in which elements of the initial trace may remain, all of which allows the animals to detect the novel spatial change. Recent data from our laboratory support this assumption, since the evocation of remote memory induced by the systemic administration of erythropoietin, i.e., 21 days after its acquisition, produces an increase in the hippocampal expression of brain-derived growth factor (BDNF) at 30 min after its induction (Almaguer-Melian W, *et al*., 2024). Moreover, memory retrieval after 21 days with the novel change of position triggers the mechanisms of memory destabilization and subsequent reconsolidation of spatial memory in the hippocampus, and the increase in the expression of BDNF could contribute to the re-stabilization and updating of the memory trace (Bekinschtein P *et al*., 2008; Bekinschtein P *et al*., 2014; Bekinschtein P *et al*., 2014; Slipczuk L *et al*., 2009).

Taking into account that a large group of DG cells is temporarily activated by BLA stimulation (Mercerón-Martínez D, *et al*., 2020) and that in contextual fear memory an engram that is formed in the DG remains silent at the memory remote time point (Kitamura T *et al*., 2017), as well as observations that the DG is important to discriminate very similar contexts (Schmidt B *et al*., 2012), we injected ZIP into DG to disrupt DG memory traces that may contribute to the maintenance of memory for precise discriminations. However, our data show that this procedure did not abolish the remote memory facilitated by amygdala stimulation. This result does not eliminate the possibility that there are memory traces not dependent on PKMζ or PKC□/ □that could contribute to maintaining spatial precision memory observed at a remote memory time point (Abraham WC *et al*., 2019; Kwapis JL *et al*., 2009; Serrano P *et al*., 2008; Tsokas P, *et al*., 2024). Future studies may clarify whether there are engram cells in the dentate gyrus of the hippocampus or CA1 that can provide a memory with sufficient details to detect the spatial change observed 21 days after learning.

The reinforcing effect of amygdala stimulation transforming a short-term memory into a remote memory is also in line with the “behavioral tagging” hypothesis. These studies, inspired by the synaptic tagging hypothesis of LTP (Frey U and Morris RG, 1997), have shown that novelty exploration close to the moment of training can produce memory consolidation (Ballarini F *et al*., 2009; Moncada D *et al*., 2015), which depends on the synthesis of new proteins (Moncada D and Viola H, 2007). Indeed, this behavioral reinforcing effect can even preserve memory for events that can interfere with its formation and promote forgetting (Almaguer-Melian W *et al*., 2012). More recently, evidence indicates that memory reconsolidation is also mediated by "behavioral tagging" process (Rabinovich Orlandi I *et al*., 2020).

Which mechanism supports amygdala-stimulation-induced facilitation of remote memory formation in the ACC? As mentioned above, when the amygdala is stimulated 15 minutes, but not 5 hours after acquisition, memory consolidation occurs (Fig. 1), and blocking aPKC action in the ACC on day 19 post-acquisition “erases” this remote memory when measured on day 21 (Fig. 3).

We propose that this is based on the synaptic tagging hypothesis (Frey U and Morris RG, 1997), which assumes that during the induction of LTP, in addition to the changes in synaptic transmission, a synaptic tag is formed. The synaptic tag then captures plasticity-related proteins, which are required for LTP maintenance beyond 3 h. We speculate that in our experiments, activity related to information processing during memory acquisition produces two dissociable events: transient plastic modifications at activated synapses, and setting of the synaptic tag at activated synapses in the hippocampus and cingulate cortex. Then amygdala stimulation 15 after the training induces synthesis of plasticity-related proteins, in particular PKMζ in the ACC. Recently, the postsynaptic scaffolding protein, KIBRA, has recently been found to act as a persistent synaptic tag for newly synthesized PKMζ in late-LTP and spatial long-term memory in hippocampus that can last weeks (Tsokas P, *et al*., 2024). Future work using antagonists of KIBRA-PKMζ interaction could test the hypothesis that KIBRA anchors PKMζ for remote memory in AAC. Other recently synthesized proteins might also be captured by the tagged synapses and in this way transform short-term memory to remote memory. For example, we previously showed that amygdala stimulation induces an increase in MAP2 and GAP43 in the prefrontal cortex and in hippocampus, strongly suggesting that structural plasticity and synaptogenesis may occur (Mercerón-Martínez D, *et al*., 2018). The persistent strengthening of pre-existing synapses maintained by PKMζ as well as the generation of new synapses could contribute to the formation of remote memory in the ACC. Indeed, previous data show that novelty exploration increases PKMζ and promotes memory consolidation in prelimbic prefrontal cortex in a behavioral-tagging process (Naseem M *et al*., 2019).

Although future studies are required to determine if the effect on remote memory of amygdala stimulation is a process that evolves over weeks or can be faster and occur in days, recent data from our laboratory suggests that it could be rapid. Relatively early expression (24 h) of a remote memory that has been reinforced by systemic administration of erythropoietin produces an increase in Arc protein expression 30 min after retrieval of the memory that is larger in the prefrontal cortex than in hippocampus (Almaguer-Melian W, *et al*., 2024). Moreover, previous studies have shown that remote memory formation can occur rapidly based on memory schemas (Tse D *et al*., 2007).

In conclusion, our data strongly indicate that activation of the amygdala shortly after the acquisition of a short-term memory can produce systems consolidation in cortical regions such as the ACC through the persistence of aPKC action. The continued action of aPKC through the autonomous activity of PKMζ maintains plasticity mechanisms by inhibiting GluA2 subunit-dependent AMPA receptor endocytosis (Migues PV *et al*., 2010; Patel H and Zamani R, 2021; Yao Y *et al*., 2008), which in turn may maintain the accumulation of the persistent synaptic tag KIBRA at activated synapses (Tsokas P, *et al*., 2024). In this way the memory persists after experience.

Overall, our results indicated the powerful modulating role of amygdala on the memory of experiences relevant to the survival of animals. We also provide data suggesting that, in natural conditions, if amygdala activation occurs close to the moment in which the information of an experience is being processed, then the additional activity of the amygdala on the memory trace can help promote the formation of remote memory of some everyday experiences; if it is not activated, memory can be lost in days or weeks. Finally, the data presented here opens a path in the search for neurological restoration based on synaptic plasticity mechanisms that can contribute to the recovery of lost functions, particularly in memory loss in aging or neurodegenerative diseases like Alzheimer’s disease.

## Acknowledgement

The authors want to thank Dr. Alain Y. García Varona and technicians Carlos Adolfo Rodríguez Pujol y Osmay Alfredo Trujillo Batista for the excellent handling and care of the experimental animals.

## Funds

The funds supporting this study were provided by CIREN and NIH funding RO1 MH115304 (T.C.S), 2R37MH057068 (T.C.S.), and R01 NS108190 (T.C.S).

## Author contributions

All authors had full access to all the data in the study and take responsibility for the integrity of the data and the accuracy of the data analysis. Study concept and design: William Almaguer-Melian, Jorge Bergado-Rosado, and Todd Sacktor. Acquisition of data: Daymara Mercerón-Martinez, William Almaguer-Melian, Laura Alacán-Ricardo, Arturo Bejerano Pina. Analysis and interpretation of data: William Almaguer-Melian, Daymara Mercerón-Martinez, Changchi Hsieh, Jorge Bergado-Rosado, and Todd Sacktor. Drafting of the manuscript: Jorge Bergado-Rosado, William Almaguer-Melian, Daymara Mercerón-Martinez, and Todd Sacktor. Statistical analysis: William Almaguer-Melian, Daymara Mercerón-Martinez, Changchi Hsieh, and Jorge Bergado- Rosado. Obtained funding: Daymara Mercerón-Martinez and Todd Sacktor. Study supervision: William Almaguer-Melian, Jorge Bergado-Rosado, and Todd Sacktor.

## References

Abraham WC, Jones OD, Glanzman DL (2019), Is plasticity of synapses the mechanism of long-term memory storage? NPJ Sci Learn 4:9.

Akirav I, Richter-Levin G (1999), Biphasic modulation of hippocampal plasticity by behavioral stress and basolateral amygdala stimulation in the rat. JNeurosci 19:10530–10535.

Akirav I, Richter-Levin G (1999), Priming stimulation in the basolateral amygdala modulates synaptic plasticity in the rat dentate gyrus. NeurosciLett 270:83–86.

Alam MJ, Kitamura T, Saitoh Y, Ohkawa N, Kondo T, Inokuchi K (2018), Adult Neurogenesis Conserves Hippocampal Memory Capacity. J Neurosci 38:6854–6863.

Almaguer-Melian W, Bergado JA, Martinez-Marti L, Duany-Machado C, Frey JU (2010), Basolateral amygdala stimulation does not recruit LTP at depotentiated synapses. Physiol Behav 101:549–553.

Almaguer-Melian W, Bergado-Rosado J, Pavón-Fuentes N, Alberti-Amador E, Mercerón-Martínez D, Frey JU (2012), Novelty exposure overcomes foot shock-induced spatial-memory impairment by processes of synaptic-tagging in rats. Proc Natl Acad Sci U S A 109:953–958.

Almaguer-Melian W, Capdevila V, Ramirez M, Vallejo A, Rosillo-Marti JC, Bergado-Rosado J (2005), Post-training stimulation of the basolateral amygdala improves spatial learning in rats with lesion of the fimbria-fornix. RestorNeurol Neurosci 23:43–50.

Almaguer-Melian W, Martinez-Marti L, Frey JU, Bergado JA (2003), The amygdala is part of the behavioural reinforcement system modulating long-term potentiation in rat hippocampus. Neuroscience 119:319–322.

Almaguer-Melian W, Mercerón-Martinez D, Alberti-Amador E, Alacán-Ricardo L, de Bardet JC, Orama-Rojo N, Vergara-Piña AE, Herrera-Estrada I, et al. (2024), Learning induces EPO/EPOr expression in memory relevant brain areas, whereas exogenously applied EPO promotes remote memory consolidation. Synapse (New York, NY) 78:e22282.

Bailey CH, Kandel ER, Harris KM (2015), Structural Components of Synaptic Plasticity and Memory Consolidation. Cold Spring HarbPerspectBiol 7.

Ballarini F, Moncada D, Martinez MC, Alen N, Viola H (2009), Behavioral tagging is a general mechanism of long-term memory formation. Proc Natl Acad Sci U S A 106:14599–14604.

Bekinschtein P, Cammarota M, Izquierdo I, Medina JH (2008), BDNF and memory formation and storage. Neuroscientist 14:147–156.

Bekinschtein P, Cammarota M, Medina JH (2014), BDNF and memory processing. Neuropharmacology 76:677–683.

Bekinschtein P, Kent BA, Oomen CA, Clemenson GD, Gage FH, Saksida LM, Bussey TJ (2014), Brain-derived neurotrophic factor interacts with adult-born immature cells in the dentate gyrus during consolidation of overlapping memories. Hippocampus 24:905–911.

Bergado JA, Frey S, Lopez J, Almaguer-Melian W, Frey JU (2007), Cholinergic afferents to the locus coeruleus and noradrenergic afferents to the medial septum mediate LTP-reinforcement in the dentate gyrus by stimulation of the amygdala. NeurobiolLearnMem 88:331–341.

Bliss TV, Lomo T (1973), Long-lasting potentiation of synaptic transmission in the dentate area of the anaesthetized rabbit following stimulation of the perforant path. JPhysiol(Lond) 232:331–356.

Campolongo P, Roozendaal B, Trezza V, Hauer D, Schelling G, McGaugh JL, Cuomo V (2009), Endocannabinoids in the rat basolateral amygdala enhance memory consolidation and enable glucocorticoid modulation of memory. Proc Natl Acad Sci U S A 106:4888–4893.

Chen Y, Barsegyan A, Nadif Kasri N, Roozendaal B (2018), Basolateral amygdala noradrenergic activity is required for enhancement of object recognition memory by histone deacetylase inhibition in the anterior insular cortex. Neuropharmacology 141:32–41.

Chen YF, Song Q, Colucci P, Maltese F, Siller-Pérez C, Prins K, McGaugh JL, Hermans EJ, et al. (2022), Basolateral amygdala activation enhances object recognition memory by inhibiting anterior insular cortex activity. Proc Natl Acad Sci U S A 119:e2203680119.

Dudai Y (2004), The neurobiology of consolidations, or, how stable is the engram? Annual review of psychology 55:51–86.

Dudai Y, Karni A, Born J (2015), The Consolidation and Transformation of Memory. Neuron 88:20–32.

Frey S, Bergado-Rosado J, Seidenbecher T, Pape HC, Frey JU (2001), Reinforcement of early long-term potentiation (early-LTP) in dentate gyrus by stimulation of the basolateral amygdala: heterosynaptic induction mechanisms of late-LTP. J Neurosci 21:3697–3703.

Frey U, Morris RG (1997), Synaptic tagging and long-term potentiation. Nature 385:533–536.

Frey U, Morris RG (1998), Weak before strong: dissociating synaptic tagging and plasticity-factor accounts of late-LTP. Neuropharmacology 37:545–552.

Hardt O, Migues PV, Hastings M, Wong J, Nader K (2010), PKMzeta maintains 1-day- and 6-day-old long-term object location but not object identity memory in dorsal hippocampus. Hippocampus 20:691–695.

Hardt O, Nadel L (2018), Systems consolidation revisited, but not revised: The promise and limits of optogenetics in the study of memory. Neuroscience letters 680:54–59.

Hatfield T, McGaugh JL (1999), Norepinephrine infused into the basolateral amygdala posttraining enhances retention in a spatial water maze task. NeurobiolLearnMem 71:232–239.

Ikegaya Y, Saito H, Abe K (1994), Attenuated hippocampal long-term potentiation in basolateral amygdala-lesioned rats. Brain Res 656:157–164.

Ikegaya Y, Saito H, Abe K (1995), High-frequency stimulation of the basolateral amygdala facilitates the induction of long-term potentiation in the dentate gyrus in vivo. NeurosciRes 22:203–207.

Ikegaya Y, Saito H, Abe K (1995), Requirement of basolateral amygdala neuron activity for the induction of long-term potentiation in the dentate gyrus in vivo. Brain Res 671:351–354.

Inman CS, Manns JR, Bijanki KR, Bass DI, Hamann S, Drane DL, Fasano RE, Kovach CK, et al. (2018), Direct electrical stimulation of the amygdala enhances declarative memory in humans. Proc Natl Acad Sci U S A 115:98–103.

Kandel E, Dudai Y, Mayford M (2014), The molecular and systems biology of memory. Cell 157:163–186.

Kitamura T, Ogawa SK, Roy DS, Okuyama T, Morrissey MD, Smith LM, Redondo RL, Tonegawa S (2017), Engrams and circuits crucial for systems consolidation of a memory. Science 356:73–78.

Krug M, L”ssner B, Ott T (1984), Anisomycin blocks the late phase of long-term potentiation in the dentate gyrus of freely moving rats. Brain ResBull 13:39–42.

Kwapis JL, Jarome TJ, Lonergan ME, Helmstetter FJ (2009), Protein kinase Mzeta maintains fear memory in the amygdala but not in the hippocampus. Behavioral neuroscience 123:844–850.

LaLumiere RT, Buen TV, McGaugh JL (2003), Post-training intra-basolateral amygdala infusions of norepinephrine enhance consolidation of memory for contextual fear conditioning. J Neurosci 23:6754–6758.

Liang KC, Juler RG, McGaugh JL (1986), Modulating effects of posttraining epinephrine on memory: involvement of the amygdala noradrenergic system. Brain Res 368:125–133.

Ling DS, Benardo LS, Serrano PA, Blace N, Kelly MT, Crary JF, Sacktor TC (2002), Protein kinase Mζ is necessary and sufficient for LTP maintenance. Nat Neurosci 5:295–296.

Matthies H (1989), In search of cellular mechanisms of memory. ProgNeurobiol 32:277–349.

McGaugh JL (2002), Memory consolidation and the amygdala: a systems perspective. Trends Neurosci 25:456–461.

McGaugh JL (2015), Consolidating memories. Annual review of psychology 66:1–24.

Mercerón-Martínez D, Almaguer-Melian W, Alberti-Amador E, Bergado JA (2018), Amygdala stimulation promotes recovery of behavioral performance in a spatial memory task and increases GAP-43 and MAP-2 in the hippocampus and prefrontal cortex of male rats. Brain Research Bulletin 142:8–17.

Mercerón-Martínez D, Almaguer-Melian W, Alberti-Amador E, Calderón-Peña R, Bergado JA (2020), Amygdala stimulation ameliorates memory impairments and promotes c-Fos activity in fimbria-fornix- lesioned rats. Synapse (New York, NY) 74:e22179.

Merceron-Martinez D, Almaguer-Melian W, Alberti-Amador E, Estupinan B, Fernandez I, Bergado JA (2016), Amygdala electrical stimulation inducing spatial memory recovery produces an increase of hippocampal bdnf and arc gene expression. Brain ResBull 124:254–61. doi: 10.1016/j.brainresbull.2016.05.017. Epub;%2016 Jun 1.:254-261.

Mercerón-Martínez D, Almaguer-Melian W, Alberti-Amador E, Estupiñán B, Fernández I, Bergado JA (2016), Amygdala electrical stimulation inducing spatial memory recovery produces an increase of hippocampal bdnf and arc gene expression. Brain Res Bull 124:254–261.

Mercerón-Martínez D, Almaguer-Melian W, Bergado JA (2022), Basolateral amygdala stimulation plus water maze training restore dentate gyrus LTP and improve spatial learning and memory. Behavioural brain research 417:113589.

Mercerón-Martínez D, Almaguer-Melian W, Serrano T, Lorigados L, Pavón N, Bergado JA (2013), Hippocampal neurotrophins after stimulation of the basolateral amygdala, and memory improvement in lesioned rats. Restorative neurology and neuroscience 31:189–197.

Migues PV, Hardt O, Wu DC, Gamache K, Sacktor TC, Wang YT, Nader K (2010), PKMζ maintains memories by regulating GluR2-dependent AMPA receptor trafficking. Nat Neurosci 13:630–634.

Moncada D, Ballarini F, Viola H (2015), Behavioral Tagging: A Translation of the Synaptic Tagging and Capture Hypothesis. Neural Plasticity:1–21.

Moncada D, Viola H (2007), Induction of Long-Term Memory by Exposure to Novelty Requires Protein Synthesis: Evidence for a Behavioral Tagging. JNeurosci 27:7476–7481.

Morena M, Colucci P, Mancini GF, De Castro V, Peloso A, Schelling G, Campolongo P (2021), Ketamine anesthesia enhances fear memory consolidation via noradrenergic activation in the basolateral amygdala. Neurobiology of learning and memory 178:107362.

Naseem M, Tabassum H, Parvez S (2019), PKM-ζ Expression Is Important in Consolidation of Memory in Prelimbic Cortex Formed by the Process of Behavioral Tagging. Neuroscience 410:305–315.

Nedaei SE, Rezayof A, Pourmotabbed A, Nasehi M, Zarrindast MR (2016), Activation of endocannabinoid system in the rat basolateral amygdala improved scopolamine-induced memory consolidation impairment. Behavioural brain research 311:183–191.

Packard MG, Cahill L, McGaugh JL (1994), Amygdala modulation of hippocampal-dependent and caudate nucleus-dependent memory processes. ProcNatlAcadSciUSA 91:8477–8481.

Parent MB, Quirarte GL, Cahill L, McGaugh JL (1995), Spared retention of inhibitory avoidance learning after posttraining amygdala lesions. BehavNeurosci 109:803–807.

Pastalkova E, Serrano P, Pinkhasova D, Wallace E, Fenton AA, Sacktor TC (2006), Storage of spatial information by the maintenance mechanism of LTP. Science 313:1141–1144.

Patel H, Zamani R (2021), The role of PKMζ in the maintenance of long-term memory: a review. Reviews in the neurosciences 32:481–494.

Paxinos G, Watson C (1998) The rat brain in stereotaxic coordinates. London: Academic Press.

Quirarte GL, Roozendaal B, McGaugh JL (1997), Glucocorticoid enhancement of memory storage involves noradrenergic activation in the basolateral amygdala. ProcNatAcadSciUSA 94:14048–14053.

Rabinovich Orlandi I, Fullio CL, Schroeder MN, Giurfa M, Ballarini F, Moncada D (2020), Behavioral tagging underlies memory reconsolidation. Proc Natl Acad Sci U S A 117:18029–18036.

Reymann K, Frey U, Matthies H (1988) A multi-phase model of synaptic long-term potentiation in hippocampal CA1 neurones: protein kinase C activation and protein synthesis are required for the maintenance of the trace. In: Synaptic plasticity in the hippocampus, vol. 1 (Haas HL, Buzs ki G, eds), pp. 126-129. Berlin, Heidelberg: Springer.

Roesler R, Parent MB, LaLumiere RT, McIntyre CK (2021), Amygdala-hippocampal interactions in synaptic plasticity and memory formation. Neurobiology of learning and memory 184:107490.

Roozendaal B, McGaugh JL (1997), Basolateral amygdala lesions block the memory-enhancing effect of glucocorticoid administration in the dorsal hippocampus of rats. EurJNeurosci 9:76–83.

Sacktor TC, Fenton AA (2012), Appropriate application of ZIP for PKMzeta inhibition, LTP reversal, and memory erasure. Hippocampus 22:645–647.

Sacktor TC, Hell JW (2017), The genetics of PKMζ and memory maintenance. 10.

Sardari M, Rezayof A, Zarrindast MR (2015), 5-HT1A receptor blockade targeting the basolateral amygdala improved stress-induced impairment of memory consolidation and retrieval in rats. Neuroscience 300:609–618.

Schmidt B, Marrone DF, Markus EJ (2012), Disambiguating the similar: the dentate gyrus and pattern separation. Behavioural brain research 226:56–65.

Seidenbecher T, Reymann KG, Balschun D (1997), A post-tetanic time window for the reinforcement of long-term potentiation by appetitive and aversive stimuli. ProcNatlAcadSciUSA 94:1494–1499.

Serrano P, Friedman EL, Kenney J, Taubenfeld SM, Zimmerman JM, Hanna J, Alberini C, Kelley AE, et al. (2008), PKMζ maintains spatial, instrumental, and classically conditioned long-term memories. PLoS Biol 6:2698–2706.

Slipczuk L, Bekinschtein P, Katche C, Cammarota M, Izquierdo I, Medina JH (2009), BDNF activates mTOR to regulate GluR1 expression required for memory formation. PLoSONE 4:e6007.

Squire LR, Genzel L, Wixted JT, Morris RG (2015), Memory consolidation. Cold Spring Harb Perspect Biol 7:a021766.

Terranova JI, Ogawa SK, Kitamura T (2019), Adult hippocampal neurogenesis for systems consolidation of memory. Behavioural brain research 372:112035.

Tonegawa S, Morrissey MD, Kitamura T (2018), The role of engram cells in the systems consolidation of memory. Nat RevNeurosci 19:485–498.

Tse D, Langston RF, Kakeyana M, Bethus I, Spooner PA, Wood ER, Witter MP, Morris RGM (2007), Schemas and Memory Consolidation. Science 316:76–82.

Tsokas P, Hsieh C, Flores-Obando RE, Bernabo M, Tcherepanov A, Hernandez AI, Thomas C, Bergold PJ, et al. (2024), KIBRA anchoring the action of PKMzeta maintains the persistence of memory. Sci Adv 10:eadl0030.

Tsokas P, Hsieh C, Yao Y, Lesburgueres E, Wallace EJ, Tcherepanov A, Jothianandan D, Hartley BR, et al. (2016), Compensation for PKMzeta in long-term potentiation and spatial long-term memory in mutant mice. eLife 5:e14846.

Wahlstrom KL, Huff ML, Emmons EB, Freeman JH, Narayanan NS, McIntyre CK, LaLumiere RT (2018), Basolateral Amygdala Inputs to the Medial Entorhinal Cortex Selectively Modulate the Consolidation of Spatial and Contextual Learning. J Neurosci 38:2698–2712.

Yao Y, Kelly MT, Sajikumar S, Serrano P, Tian D, Bergold PJ, Frey JU, Sacktor TC (2008), PKMζ maintains late long-term potentiation by N-ethylmaleimide-sensitive factor/GluR2-dependent trafficking of postsynaptic AMPA receptors. J Neurosci 28:7820–7827.

